# Making the invisible enemy visible

**DOI:** 10.1101/2020.10.07.307546

**Authors:** Tristan Croll, Kay Diederichs, Florens Fischer, Cameron Fyfe, Yunyun Gao, Sam Horrell, Agnel Praveen Joseph, Luise Kandler, Oliver Kippes, Ferdinand Kirsten, Konstantin Müller, Kristoper Nolte, Alex Payne, Matt Reeves, Jane Richardson, Gianluca Santoni, Sabrina Stäb, Dale Tronrud, Lea von Soosten, Christopher Williams, Andrea Thorn

**Affiliations:** CIMR, University of Cambridge, UK; University of Constance, Germany; HARBOR, Universität Hamburg, Germany; Paris, France; RVZ, University of Würzburg, Germany; Diamond Light Source, UK; Science and Technology Facilities Council, UK; Memorial Sloan Kettering Cancer Center, USA; Duke University, USA; European Synchrotron Radiation Facility, France; Oregon, USA

## Abstract

During the COVID-19 pandemic, structural biologists rushed to solve the structures of the 28 proteins encoded by the SARS-CoV-2 genome in order to understand the viral life cycle and enable structure-based drug design. In addition to the 204 previously solved structures from SARS-CoV-1, 548 structures covering 16 of the SARS-CoV-2 viral proteins have been released in a span of only 6 months. These structural models serve as the basis for research to understand how the virus hijacks human cells, for structure-based drug design, and to aid in the development of vaccines. However, errors often occur in even the most careful structure determination - and may be even more common among these structures, which were solved quickly and under immense pressure.

The Coronavirus Structural Task Force has responded to this challenge by rapidly categorizing, evaluating and reviewing all of these experimental protein structures in order to help downstream users and original authors. In addition, the Task Force provided improved models for key structures online, which have been used by Folding@Home, OpenPandemics, the EU JEDI COVID-19 challenge and others.

## Introduction

The coronavirus SARS-CoV-2 has a single-stranded RNA genome encoding 28 proteins. These enable SARS-CoV-2 to infect, replicate, and suppress the immune system of its host. For example, the characteristic spikes that protrude from its envelope and allow it to bind to host cells are a trimer of the surface glycoprotein (Fig. 1). Knowing the atomic structures of these macromolecules is vital for understanding the life cycle of the virus and helping design specific pharmaceutical compounds that can inhibit their functions, with the goal of stopping the cycle of infection.

**Fig. 1.**
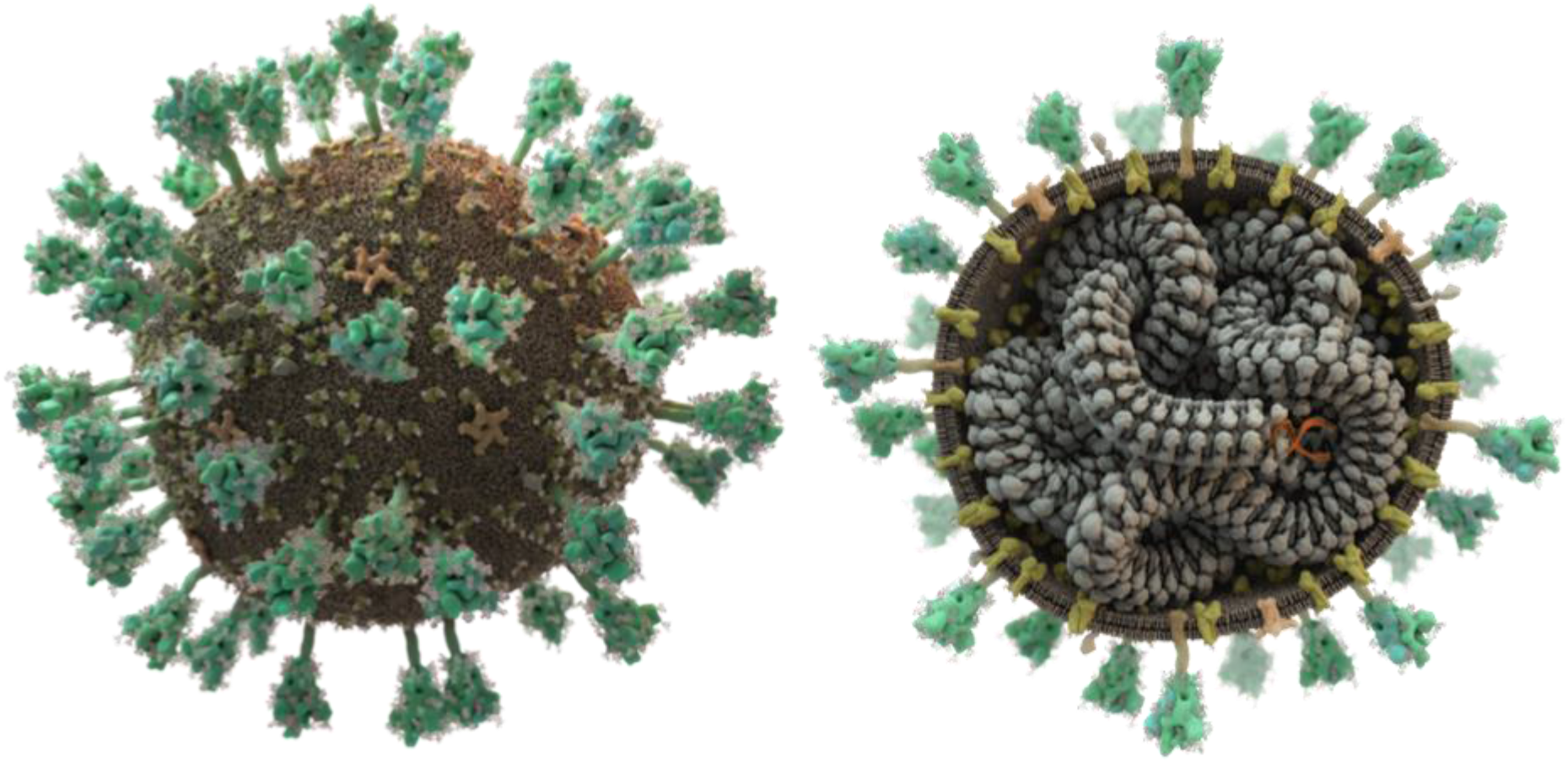
SARS-CoV-2 displays spike proteins (green) on its surface that recognize and bind to host cells; its lipid bilayer membrane also contains additional embedded membrane and envelope proteins (yellow and beige, respectively). The single-stranded RNA (orange) is intertwined in a helical fashion with the nucleocapsid (grey). This figure only shows only the transport form of the virus: once a cell is infected, additional viral proteins encoded by the viral RNA are produced that hijack the host cell in order to produce new virus particles. (Picture: Thomas Splettstöβer /scistyle.com)

When the COVID-19 pandemic hit in the beginning of this year the structural biology community swung into action very efficiently and is now strongly engaged to determine the atomic structures of the viral macromolecules as fast as possible^1^. All structure determination methods require interpretation of the measured and processed data with a structural model and cannot be fully automated. The resulting structures are freely and publicly available in the World Wide Protein Data Bank (wwPDB), structural biology’s archive of record^2^.

Unfortunately, the fit between model and data is never perfect, and errors from measurement, post-processing and modelling are a given. Structures solved in a hurry to address a pressing medical and societal need may be even more prone to mistakes. These small faults can have severe consequences – in particular, in structure-based drug discovery, structural bioinformatics, and computational chemistry; all of which are being employed in current SARS-CoV-2 research across the world. As of the writing of this publication, 524 macromolecular structures from SARS-CoV-1 and SARS-CoV-2 have been deposited, covering parts of 16 of the 28 viral proteins. In this time of crisis, it is vital to ensure that the structural data made available to the wider research community are the best they can be in every regard.

### Pushing the methods to the limit

The wwPDB^3^ is an invaluable tool, but it is a near-static archive, where a released structure can only be updated by the original depositors. After any associated papers are published, there is often little or no motivation to make corrections.99% of structure downloads from the PDB are not by those who determine the structures, but scientists from other fields^2^. Consequently, errors can be misinterpreted as biologically and pharmaceutically relevant information and can cause a large waste of resources and time down-the-line.

As methods developers for structure determination and validation, we are expert users of our own software tools and are ideally placed to help in this unprecedented situation. We joined forces to assess and, where necessary, improve upon the published macromolecular structures from SARS-CoV-1 and SARS-CoV-2. In cases where we could significantly improve the molecular models, we offered them back to the original authors and the scientific community. After we began this validation effort we were approached by researchers in *in-silico* drug screening, Folding@Home^4,5^, OpenPandemics^6^, and the EU Joint European Disruptive Initiative (JEDI)^7^. These initiatives needed the best structures they could get for studies of the virus and had already lost much computing time and resources working with suboptimal models.

### Automatic evaluation

All macromolecular structures from SARS-CoV-1 and SARS-CoV-2 in the wwPDB are downloaded into our repository and assessed automatically in the first 24 hours after release. For crystallographic and Cryo-EM structures, we check the quality of the deposited merged data, and how well the model fits these data. All structures are checked according to chemical prior knowledge.

#### Evaluation specific to crystallographic data and structure solutions

As crystal structures make up 73% of reported SARS-CoV-1 and SARS-CoV-2 structures, these are evaluated most thoroughly for pathologies such as twinning, multiple lattice diffraction, ice crystal diffraction contamination, incompleteness resulting from a poorly chosen collection strategy, limited diffraction (at low and high resolution boundaries) or radiation-induced damage which renders some reflections inaccurate. These issues cannot be resolved after data collection is complete, but explicitly taking them into account during data processing and structure solution can yield a better structural model. However, finding such problems from deposited structure factors can be difficult. Raw data allow a much more complete analysis of the experiment, including anisotropic diffraction limits, systematic errors from the measurement setup or strategy, as well as errors arising from data processing such as incorrect lattice symmetry or inclusion of diffraction spots obscured by the beam stop. If raw data are available, data can be re-processed, and these problems may be resolved. However, raw data are not deposited in the wwPDB, or required for publication, and can be difficult to obtain. Their absence in the public record is detrimental for re-analysis and evaluation. To support evaluation of the entire structure determination from start-to-finish, we invited authors to send us their raw experimental data and offered to deposit them in a public repository, such as SBGrid^8^ or proteindiffraction.org^9^.

To analyse crystallographic data for twinning, completeness, and overall diffraction quality, we used phenix.xtriage^10^; to identify ice rings and additional pathologies, we used AUSPEX^11^ (see Fig. 2F). The overall completeness of most datasets is satisfactory, with only 7 out of 547 datasets below 80%. All data sets have an acceptable signal-to-noise. Ice rings were detected in 61 datasets and problems with the beam stop masking in 54; 41 crystal structures were indicated as potentially resulting from twinned crystals.

**Fig. 2.**
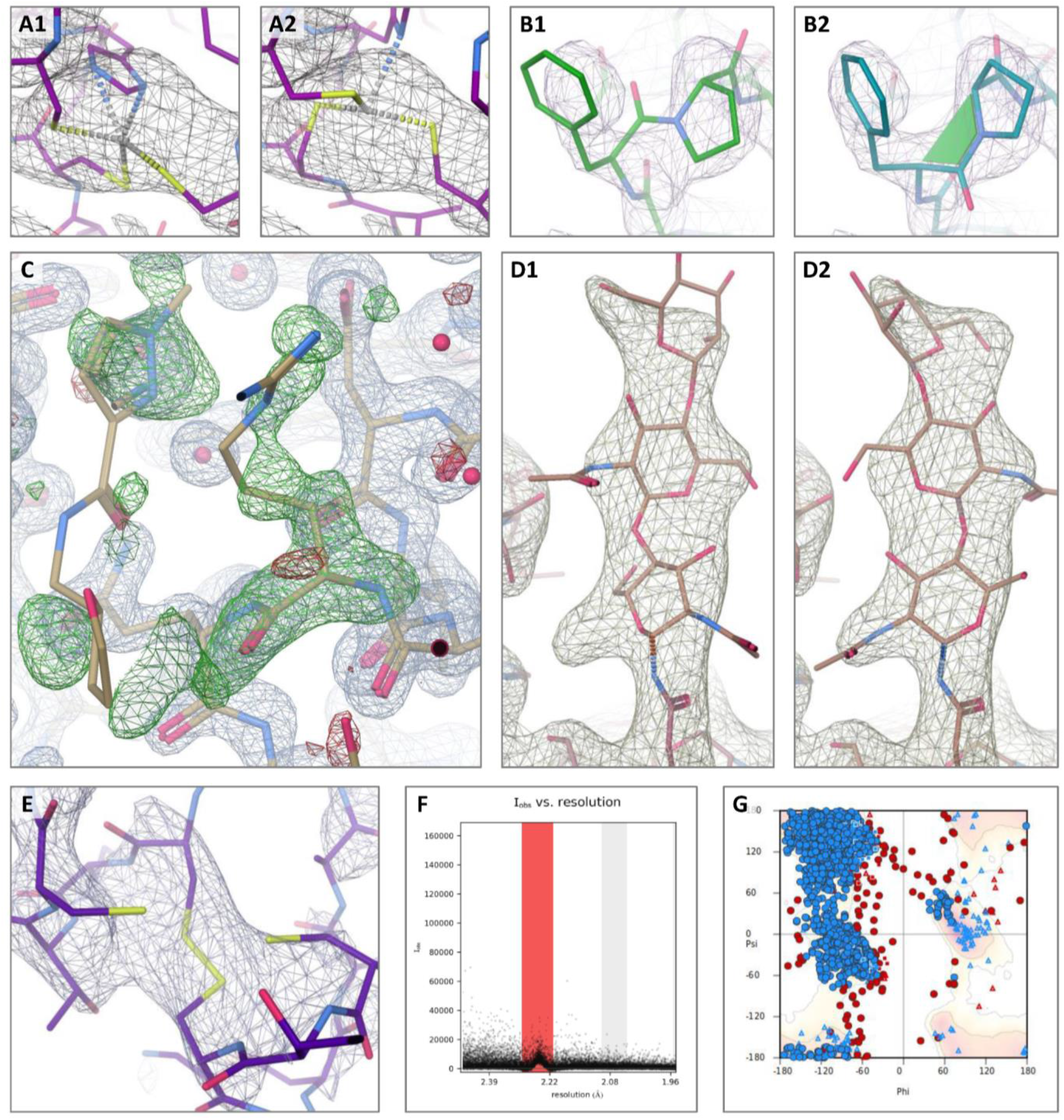
Indicators for improvement. All pictures except F are screenshots from Coot 0.9.9-pre-release. Residual density and reconstructions maps in blue; difference density in red and green. **A1.** SARS coronavirus nsp14-nsp10 (PDB 5c8t) histidine zinc coordination site (B603) with residual density contour level 0.445, rmsd 0.150. **A2.** Histidine has been swapped in ISOLDE, leading to tetrahedral coordination of Zn^2+^, followed by PDB-REDO refinement with manually added links. **B1./B2.** Proline A505 is modelled as trans in RdRp complex (PDB 7bv2, left) but density indicates a cis main chain conformation (right). After we contacted the original authors, this was changed. **C.** Residue A165 (right) with 0.44 occupancy instead of 1.00 near potential inhibitor (left) in SARS-CoV-2 main protease (PDB 5rfa); this happened frequently in entries from the XChem/COVID-Moonshot project. Residual map contour level 0.54, rmsd 0.319, difference density at contour level 0.35, rmsd 0.114. **D1.** SARS-CoV-2 spike receptor binding domain complexed with human ACE2 (PDB 6vw1): an entire N-linked glycan is flipped approximately 180° around the N-glycosidic bond. After we contacted the original authors, this entry was revised - see **D2.** Correction improves the density fit of the sugar chain (right). Residual map at contour level 0.311 rmsd 0.265. **E.** Disulphide bond A226-A189 in papain-like protease (PDB 6w9c) with electron density at contour level 0.214 rmsd 0.136. While the density map does not indicate a zinc, it is a zinc finger domain; the other NCS copies have a zinc coordinated here and the other two cysteines are uncoordinated. **F.** AUSPEX plot of SARS-CoV main protease (PDB 2hob); ice rings are reflected by a bias in the intensity distribution (red). **G.** Ramachandran plot for SARS-CoV nsp10/nsp14 dynamic complex (PDB entry 5nfy).

A general indication of how well the atomic model fits the measurement data can be obtained from the R factors: While only two structures in the repository present an alarmingly high *R*_free_ value, above 35%, this does not indicate the absence of modelling problems in the others. We supply PDB-REDO^12^ re-refinements which generally improve *R*_free_, but the resulting models should not be viewed as “more correct” purely on basis of a lower R value, particularly at lower resolution where the relationship between R factors and model quality breaks down^13^. Critical inspection of the model details in the density remains necessary (while being cautious to avoid expectation and model bias). Large *R*_free_ drops indicated major issues with the deposited PDB entry; this was especially true for older SARS-CoV-1 structures.

#### Evaluation specific to structures from single-particle Cryo-EM

Cryo-EM structures make up 23% of reported SARS-CoV-1 and SARS-CoV-2 structures. As with crystallographic structures, raw data are not available from the wwPDB, but deposition into EMPIAR^14^ is increasingly common. The reconstructed 3D map deposited in the EMDB^15^ allows calculation of the fit between model and map in the form of a Fourier Shell Correlation (FSC) to assess agreement between features at different resolutions. For a well-fitted model, the cryo-EM map resolution (defined as where the FSC between two independently calculated half-maps drops below 0.143) roughly corresponds to a model-map FSC of 0.5. To calculate FSCs, we utilized the CCP-EM^16^ model validation task which uses REFMAC5^17^ and additionally calculates real-space Cross-Correlation Coefficient (CCC), Mutual Information (MI) and Segment Manders’ Overlap Coefficient (SMOC)^18^. While MI is a single value score to evaluate how well model and map agree, the SMOC score evaluates the fit of each modelled residue individually and can help to find regions where errors occur in the model in relation to the map. Z-scores highlight residues with a low score relative to their neighbors and point to potential misfits.

Of 120 structures, 20 structures had an average model-map FSC below 0.4 and 21 had a MI score below 0.4, indicating poor overall agreement between map and model, and potential for further improvement. The SMOC score of 28 structures indicated that more than 10% of the residues fit poorly with the map. These models could potentially be improved to better fit the map reconstructed from the experimental data. In addition to the above validation, we run Haruspex^19^, a neural network trained to recognize secondary structure elements and RNA/DNA in Cryo-EM maps, as visual guidance for structure evaluation.

#### Evaluation of the structural models based on prior knowledge

In order to evaluate the model quality with respect to chemical prior knowledge we run MolProbity^20,21^, which checks covalent geometry, conformational parameters of protein and RNA, and steric clashes. However, some of these traditional indicators of model quality are often used as additional restraints during refinement, which invalidates their use as quality metrics to a certain degree – we therefore used the MolProbity CaBLAM score^22^, which can pinpoint local errors at 3-4 Å resolution, even if traditional criteria have been used as restraints during refinement. CaBLAM indicated that 131 of the structures have many uncommon local backbone conformations (> 2% outliers); sometimes this was already evident from Ramachandran plots (see Fig. 2G). During the Corona crisis the MolProbity webservice has been pushed to the limit of its capacity, as many different drug developers screen the same coronavirus structures many times.

We developed a custom MolProbity pipeline which makes the validation results for each coronavirus structure available online to decrease the workload on the webservice. In addition to this, the sequence of each structure is checked against the known genome; this highlighted misidentified residues in 23 structures. Modelling errors in general can be corrected, but mostly need manual intervention (see below).

#### Online availability and updates

Every Wednesday, when new PDB structures are released, our automatic pipeline identifies new coronavirus structures and assesses the quality of both models and experimental results. This assessment, along with the original structures, are then immediately made available from our online repository which is accessible via our website insidecorona.net. To facilitate access and provide an overview of structures, we supply a summary, an SQL database of key statistics and quality indicators, as well as the individual results.

### Manual evaluation

The structural biology community has achieved a high level of automation in recent years. However, due to the complexity of interpreting low-quality maps that have poor fit between experimental data and structural models, the process of structure-solution still requires guidance and interpretation by researchers, as it has not yet been fully automated. Experienced human inspection residue-by-residue remains the best way to judge the quality of a structure, highlighting the continuing need for expert structure solvers. Given the flood of new SARS-CoV-2 structures, we have selected representative structures from each SARS-CoV-2 protein as well as those of particular interest as reference structures in drug development. Certain errors were surprisingly common, such as peptide bond flips (see Fig 2, B1 and B2), rotamer outliers, occupancy problems (see Fig. 2C) and mis-identification of small molecules/ions, such as water as magnesium, chloride as zinc. Zinc plays an important role in many viral infections, and is coordinated by many of the SARS-CoV-2 protein structures. We found a large number of Cys-Zn sites to be mismodelled, with the zinc ion missing or pushed out of density, and/or erroneous disulphide bonds between the coordinating cysteine residues (see Fig 2A1, A2, E). Many coronavirus proteins are glycosylated at surface asparagine residues. In many structures containing glycans the sugars were flipped approximately 180 degrees from their correct orientation around the N-glycosidic bond (see Fig. 2, D1 and D2). This could be avoided using tools such as Privateer^23^ and the automated carbohydrate building tool in Coot^24^.

Undeniably, deviation from prior knowledge is not universally an error. If the experimental data strongly support this deviation, it may not only be acceptable but can be an important and functionally relevant feature, as exemplified by the strained geometries often found at catalytic sites.

Of the structures we checked manually, we were able to significantly improve 31 (available from insidecorona.net) in terms of model quality, data quality, or both. Here we give two examples to illustrate the importance of careful inspection of experimental data and resulting models.

#### Example 1: Papain-like protease

Upon SARS-CoV-2 infection, a long polypeptide chain is produced and then cleaved into 16 functional proteins, the non-structural proteins (NSPs)^25^ which are essential for replication of the virus. NSP3 is a large protein consisting of 1945 amino acid residues in 15 segments with a variety of functions. One of these is the papain-like protease domain which cleaves the first five NSPs from the polypeptide chain^25^ in an essential step for viral replication, making it an important drug target^26^.

The first SARS-CoV-2 structure of this domain (PDB 6W9C) was released 1^st^ April 2020 and was immediately used in drug design efforts around the world. The overall completeness of the measured data, however, was only 57%. Why were all of the data not recorded?

Examination of the raw data, available from proteindiffraction.org^9^, revealed that the data had been measured with a very high X-ray intensity, leading to a rapid loss of higher resolution data due to radiation damage. This is not possible to deduce from the information deposited in the PDB, underlining the importance of the availability of raw data. The crystals were small thin plates; thus, dose distribution is crucial in order to achieve a serviceable experimental dataset. We estimated the diffraction weighted average dose applied to this crystal is 5.5MGy, an acceptable albeit high value. However, the maximum applied dose is estimated as 21MGy, which surely would have destroyed part of the crystal^27^. A second issue apparent from the raw data is that the data were collected with a 30° sample rotation followed by a 60° rotation starting from the same position, covering the first 30° twice. This increased the damage while yielding little additional information. The angular range per image of 0.5°, was also surprisingly wide for a pixel array detector.

The crystal has 3-fold non-crystallographic symmetry (NCS) with the monomers containing a functionally important zinc ion bound by four cysteines. The Zn-sites have poor density, which may be the result of radiation damage, and the 3 subunits have differing and incomplete Zn-site models. Only two monomers have zinc ions modelled, but the bond lengths between the four sulphur atoms and the zinc vary from 2.4 Å to 2.7 Å and the C^ß^-SG-Zn angles between 70° and 132°.The bond lengths between Cys and Zn should in fact be approximately 2.3 Å and the angles about 107°^28^. The third site is modelled as a disulphide bond and two free cysteines (Fig. 2).

We reprocessed the images using XDS^29^, omitting the final 10° of the first sweep and final 20° of the second where the radiation damage most severely affected the data. Staraniso^30^ was used to apply an anisotropic data cutoff. This careful manual intervention improved the resolution from 2.7 to 2.6 Å with better data quality overall, however, the revised overall ellipsoidal completeness was only 44.5%. The deposited model did have one of the zinc sites modelled as a disulphide bond. The other two lateral zinc sites had been refined without restraints to coordination geometry, resulting in varying S-Zn bond distances and instead of the expected 107° values. Adding zincs to all sites, restraining the bond lengths and angles to the expected values, using NCS restraints and an overall higher weighting of ideal geometry, and some remodeling of side chains and water molecules improved the electron density maps and lowered the R values by 4% to 20.2%/25.4% at 2.6 Å resolution.

This example shows the interconnected importance of data collection strategy, data processing and model building. Even though the crystal was radiation damaged, by reprocessing the data, modifying refinement to include stronger restraints, and taking full advantage of the non-crystallographic symmetry, this structure could be drastically improved. A structure of the C111S mutant of the same protein (PDB 6WRH) came out a month later, in which the zinc sites were clearly resolved in all subunits. By this time, however, 6W9C had already been widely used in *in silico* drug design: for example, 20% of participants in the EU JEDI COVID-19 challenge have used this model. The availability of a better structure a month earlier would not only have increased their chances of success but also saved much computing and person hours in computer aided drug development.

#### Example 2: RNA polymerase complex

SARS-CoV-2 replicates its single-stranded RNA genome with a macromolecular complex of RNA-dependent RNA polymerase (NSP12, RdRp), NSP7 and NSP8^31^. The first structure of SARS-CoV-1 RNA polymerase (PDB 6NUS) was solved in 2019 by Cryo-EM. In this structure, a loop close to the C-terminus (residues 892-906) was not resolved in the reconstruction map and hence not modelled. Following this loop, the polymerase has an irregular helix followed by a flexible tail. Density for this helix was poorly resolved, and its short length and lack of information from the surrounding loops led to difficulty in assigning the identity of the amino acids at each site. Nevertheless, the overall validation statistics provided by the wwPDB for this model appear to be exceptionally good. We inspected one of the first available EM structures of the equivalent SARS-CoV-2 complex (7BTF) using ISOLDE^13^. To our surprise, the higher resolution in the C-terminal region made it clear that the C-terminal helix model was completely incorrect, with the assigned sequence in fact being nine residues upstream of the correct residues for this site (see Fig. 3). This error was present in all the structures of this complex from both SARS-CoV-1 and SARS-CoV-2, propagated due to the standard practice of using the previous (incorrect) model as a starting point for each subsequent structure. For each affected structure we immediately contacted the original authors. 14 of the 18 RNA polymerase complexes in the wwPDB now have the corrected sequence alignment at the C-terminus, and also include many of our other changes described below. These PDB re-versioned corrections allow modelling efforts for drugs against SARS-CoV-2 RNA polymerase to start from a much better model. Notably, the authors of a later cryo-EM structure of the RNA polymerase/RNA complex (PDB entry 6YYT) used one of our corrected models as the starting point for their new model^32^.

**Fig. 3.**
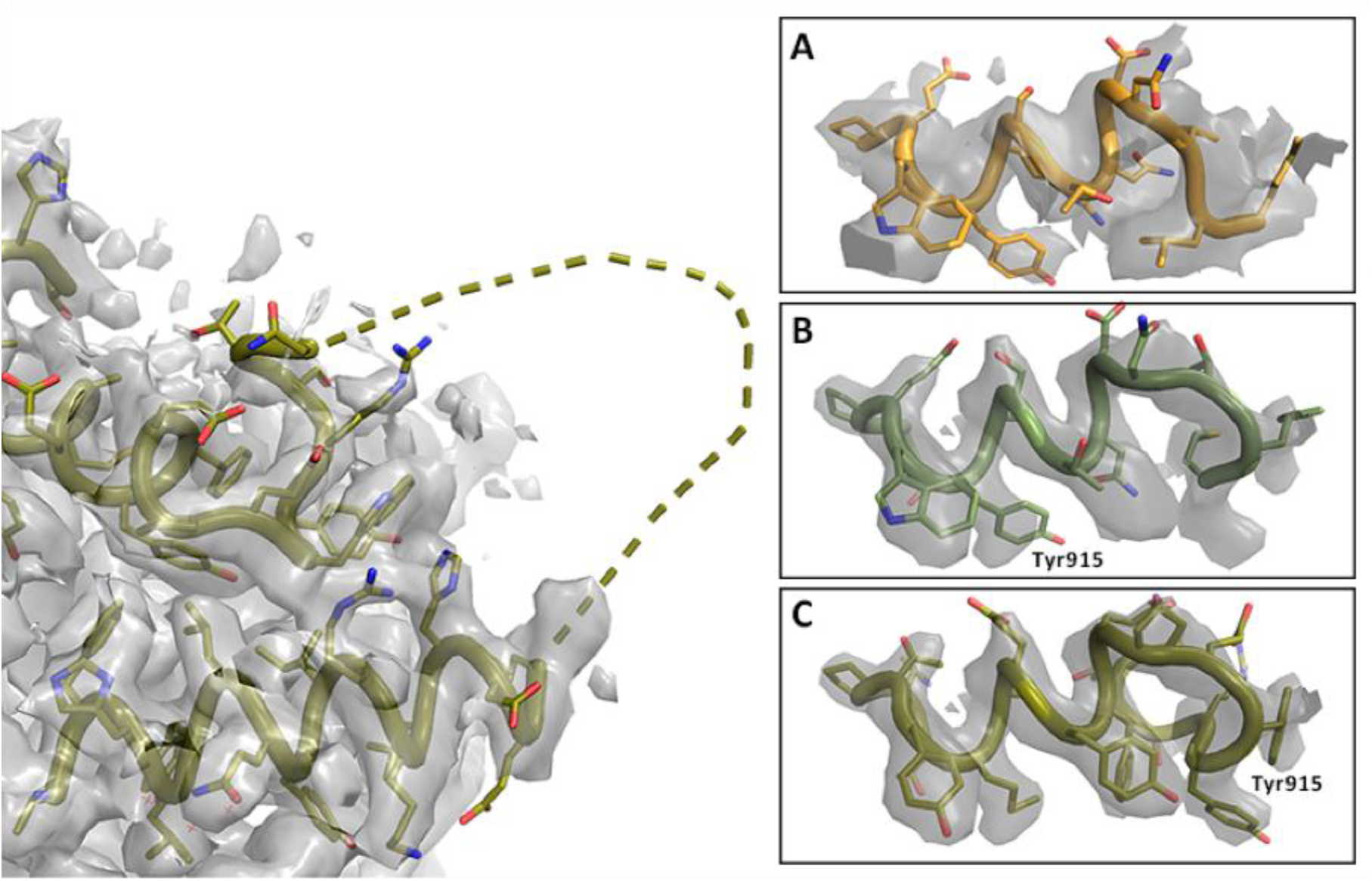
Registry shift in C-terminus of RNA Polymerase. **Left:** Overview with missing loop shown as dashed line (PDB entry 7BV2); map at 2.4σ. **Right:** Details of C-terminal helix at 5σ. **A.** Lower resolution map and model PDB 6NUS. Judging the side chain fit is difficult. **B.** Higher resolution map and model 7BV2 as deposited; the side chain fit is suboptimal. **C.** Amended 7BV2 structure; the side chains now fit the density. The register shift is indicated by Tyr915.

The structure of SARS-CoV-2 RNA polymerase in pre-translocation state and bound to template-primer RNA and Remdesivir (PDB entry 7BV2^33^) represents a useful basis for the rational design of other nucleoside triphosphate (NTP) analogues^34^. However, we found that this structure has several issues in addition to the register shift described above, which may mislead downstream users. Of greatest concern for subsequent *in-silico* docking and drug design studies is that the three magnesium ions and pyrophosphate modelled in the active site are poorly supported by the experimental data and by local geometry. We rebuilt this structure, correcting the residue shift, deleting the active site magnesium ions and pyrophosphate (replacing one magnesium with a more chemically sensible water), and improving the conformations of three RNA residues close to the Remdesivir binding site (including an adenosine base (T18) modelled “backwards”). We also fixed a number of peptides (see Fig 1, B1 and B2), added several residues and water molecules with good density and geometry, and altered two proline residues that had been erroneously flipped from *cis* to *trans*. Further methodological detail for this and other examples is available in press^35^. Interestingly, by the standard wwPDB metrics (clashscore, Ramachandran and rotamer outliers) the new model is almost indistinguishable from the old – an object lesson in the importance of human “sanity-checking” of models prior to final deposition. It should be noted, however, that most of the backwards peptide bonds and erroneous *trans* prolines were detected by the Molprobity CaBLAM^22^ feature. Our remodelled structure offers a valuable basis for future studies, such as *in-silico* docking and drug design targeting SARS-CoV-2 RdRp^36^, as well as for computational modelling and simulation to investigate the molecular mechanism of viral replication^34,37,38^.

## Supplying context

Many specialists in structural biology and *in silico* design are tackling SARS-CoV-2 research, but lack familiarity with wider coronavirus research. In addition to structure evaluation and improvement, our homepage insidecorona.net supplies literature reviews centered on the structural aspects of the viral life cycle, host interaction partners, illustrations, and selecting the best starting models for *in silico* projects. Furthermore, we added SARS-CoV-2 proteins to Proteopedia^39^ and MolSSI^40,41^, as well as a 3D-Bionotes^22^ deep-link into our database. Finally, as SARS-CoV-2 has had a large impact on the whole world, we have tried to make related research accessible to the general public. We have a number of posts aimed at non-scientists, have live streamed data processing on Twitch, and produced an accurate 3D printed model of SARS-CoV-2 based on deposited structures along with the files and instructions necessary to print these models for outreach activities.

## Summary

In the last five months, we have done a weekly automatic post-analysis as well as a manual re-processing and re-modelling of representative structures from each of the 16 structurally known macromolecules of SARS-CoV-1 or SARS-CoV-2. In this global crisis, where the community aims to get structures out as fast as possible, we aim to ensure that structure interpretations available to downstream users are as reliable as possible. We provide these results as a free resource to the community in order to aid the hunt for a vaccine or anti-viral treatment. Our results are constantly updated and can be found online at **insidecorona.net.** New contributors to this effort are very welcome.

## Outlook

In the last 40 years, structural biology has become highly automated, and methods have advanced to the point that it is now feasible to solve a new structure from start to finish in a matter of weeks with little specialist knowledge. The extremely rapid and timely solution of these structures is a remarkable achievement during this crisis and, despite some shortcomings, these structures have enabled downstream work on therapeutics to rapidly progress. The downside is that errors at all points during a structure determination are not only common, but can also remain undetected. Unfortunately, no individual researcher is fully conversant with all the details of structure determination, protein and nucleic acid structure, chemical properties of interacting groups, catalytic mechanisms, and viral life cycle. The result is that the first draft of a molecular model often contains errors like the ones pointed out above. While any molecular model could benefit from an examination by multiple experts, during this time it is important to bring such inspection to Coronavirus-related structures as quickly as possible. Raw data would allow a more complete inspection of the structure solution and a possibility to propose updates to the original authors directly, or to deposit derivative models in the wwPDB that may enhance the usefulness of these structures later.

We believe that, as a community, we need to change how we all see, address and document errors in structures to achieve the best possible structures from our experiments. We are scientists: *In the end, truth should always win*.

## Acknowledgements

This work was supported by the German Federal Ministry of Education and Research [grant no. 05K19WWA], Deutsche Forschungsgemeinschaft [project TH2135/2-1], the Wellcome Trust [grants 208398/Z/17/Z and 209407/Z/17/Z], and the US National Institutes of Health [grant R35 GM131883]. It would not have been possible without exchange, discussions and support from the computational and experimental structural biology community; particularly Lu Zhang, John Chodera, Stefano Forli, Thomas Hermanns, Paul Emsley, Clemens Vonrhein, Iris Young, James Fraser and Arwen Pearson. We would also like to thank to Holger Theymann, Nicole Dörfel and Thomas Splettstößer for web design and visualization of our work. Lastly, we are grateful to Elisa Bandello, Pairoh Seeliger & Florian Platzmann for their support.

## Literature

1. Baker, E. N. Visualizing an unseen enemy; mobilizing structural biology to counter COVID-19. Acta Cryst D 76, 311–312 (2020).

2. Burley, S. K. et al. RCSB Protein Data Bank: Sustaining a living digital data resource that enables breakthroughs in scientific research and biomedical education. Protein Science: A Publication of the Protein Society 27, 316 (2018).

3. Berman, H., Henrick, K. & Nakamura, H. Announcing the worldwide Protein Data Bank. NSMB 10, 980 (2003).

4. Shirts, M. & Pande, V. S. Screen Savers of the World Unite! Science 290, 1903–1904 (2000).

5. Zimmerman, M. I. et al. Citizen Scientists Create an Exascale Computer to Combat COVID-19. bioRxiv 2020.06.27.175430 (2020) doi:10.1101/2020.06.27.175430.

6. OpenPandemics - COVID-19 | Research | World Community Grid. https://www.worldcommunitygrid.org/research/opn1/overview.do.

7. JEDI COVID-19 Grand Challenge. https://www.covid19.jedi.group.

8. Morin, A. et al. Cutting edge: Collaboration gets the most out of software. eLife 2, (2013).

9. Grabowski, M. et al. A public database of macromolecular diffraction experiments. Acta Cryst D 72, 1181–1193 (2016).

10. Zwart, P. H., Grosse-Kunstleve, R. W. & Adams, P. D. Xtriage and Fest: automatic assessment of X-ray data and substructure structure factor estimation. 9.

11. Thorn, A. et al. AUSPEX: a graphical tool for X-ray diffraction data analysis. Acta Cryst D 73, 729–737 (2017).

12. Rp, J., F, L., Gn, M. & A, P. The PDB_REDO Server for Macromolecular Structure Model Optimization. IUCrJ vol. 1 https://pubmed.ncbi.nlm.nih.gov/25075342/ (2014).

13. Croll, T. I. ISOLDE: a physically realistic environment for model building into low-resolution electron-density maps. Acta Cryst D 74, 519–530 (2018).

14. Iudin, A., Korir, P. K., Salavert-Torres, J., Kleywegt, G. J. & Patwardhan, A. EMPIAR: a public archive for raw electron microscopy image data. Nat Methods 13, 387–388 (2016).

15. Lawson, C. L. et al. EMDataBank.org: unified data resource for CryoEM. Nucleic Acids Research 39, D456 (2011).

16. Burnley, T., Palmer, C. M. & Winn, M. Recent developments in the CCP-EM software suite. Acta Cryst D 73, 469–477 (2017).

17. Murshudov, G. N. et al. REFMAC5 for the refinement of macromolecular crystal structures. Acta Cryst D 67, 355–367 (2011).

18. Ap, J. et al. Refinement of atomic models in high resolution EM reconstructions using Flex-EM and local assessment. Methods (San Diego, Calif.) vol. 100 https://pubmed.ncbi.nlm.nih.gov/26988127/ (2016).

19. Mostosi, P., Schindelin, H., Kollmannsberger, P. & Thorn, A. Haruspex: A Neural Network for the Automatic Identification of Oligonucleotides and Protein Secondary Structure in Cryo-Electron Microscopy Maps. Angewandte Chemie International Edition (2020) doi:10.1002/anie.202000421.

20. Chen, V. B. et al. MolProbity: all-atom structure validation for macromolecular crystallography. Acta Cryst D 66, 12–21 (2010).

21. Williams, C. J. et al. MolProbity: More and better reference data for improved all-atom structure validation. Protein Science: A Publication of the Protein Society 27, 293 (2018).

22. Prisant, M. G., Williams, C. J., Chen, V. B., Richardson, J. S. & Richardson, D. C. New tools in MolProbity validation: CaBLAM for CryoEM backbone, UnDowser to rethink “waters,” and NGL Viewer to recapture online 3D graphics. Protein Science 29, 315–329 (2020).

23. Agirre, J. et al. Privateer: software for the conformational validation of carbohydrate structures. Nat Struct Mol Biol 22, 833–834 (2015).

24. Emsley, P. & Crispin, M. Structural analysis of glycoproteins: building N-linked glycans with Coot. Acta Cryst D 74, 256–263 (2018).

25. Lei, J., Kusov, Y. & Hilgenfeld, R. Nsp3 of coronaviruses: Structures and functions of a large multi-domain protein. Antiviral Research 149, 58 (2018).

26. Harcourt, B. H. et al. Identification of Severe Acute Respiratory Syndrome Coronavirus Replicase Products and Characterization of Papain-Like Protease Activity. Journal of Virology 78, 13600 (2004).

27. Bury, C. S., Brooks-Bartlett, J. C., Walsh, S. P. & Garman, E. F. Estimate your dose: RADDOSE-3D. Protein Science 27, 217–228 (2018).

28. Laitaoja, M., Valjakka, J. & Jänis, J. Zinc Coordination Spheres in Protein Structures. https://pubs.acs.org/doi/abs/10.1021/ic401072d (2013) doi:10.1021/ic401072d.

29. Kabsch, W. XDS. Acta Cryst D 66, 125–132 (2010).

30. Tickle, I. J. et al. STARANISO. (Global Phasing Ltd., 2018).

31. Smith, E. C. & Denison, M. R. Implications of altered replication fidelity on the evolution and pathogenesis of coronaviruses. Current Opinion in Virology 2, 519 (2012).

32. Hillen, H. S. et al. Structure of replicating SARS-CoV-2 polymerase. Nature 1–6 (2020) doi:10.1038/s41586-020-2368-8.

33. Yin, W. et al. Structural basis for inhibition of the RNA-dependent RNA polymerase from SARS-CoV-2 by remdesivir. Science 368, 1499–1504 (2020).

34. Zhang, L. et al. Role of 1’-Ribose Cyano Substitution for Remdesivir to Effectively Inhibit both Nucleotide Addition and Proofreading in SARS-CoV-2 Viral RNA Replication. bioRxiv 2020.04.27.063859 (2020) doi:10.1101/2020.04.27.063859.

35. Croll, T., Williams, C., Chen, V. B., Richardson, D. C. & Richardson, J. S. Improving SARS-CoV-2 structures: Peer review by early coordinate release. Biophysical Journal [in press] (2020).

36. Zhang, L. & Zhou, R. Structural Basis of the Potential Binding Mechanism of Remdesivir to SARS-CoV-2 RNA-Dependent RNA Polymerase. The Journal of Physical Chemistry B (2020) doi:10.1021/acs.jpcb.0c04198.

37. Barakat, K., Ahmed, M., Tabana, Y. & Ha, M. A “Deep Dive” into the SARS-Cov-2 Polymerase Assembly: Identifying Novel Allosteric Sites and Analyzing the Hydrogen Bond Networks and Correlated Dynamics. bioRxiv 2020.06.02.130849 (2020) doi:10.1101/2020.06.02.130849.

38. Shannon, A. et al. Remdesivir and SARS-CoV-2: Structural requirements at both nsp12 RdRp and nsp14 Exonuclease active-sites. Antiviral Research 178, 104793 (2020).

39. Proteopedia: A status report on the collaborative, 3D web-encyclopedia of proteins and other biomolecules. Journal of Structural Biology 175, 244–252 (2011).

40. Krylov, A. et al. Perspective: Computational chemistry software and its advancement as illustrated through three grand challenge cases for molecular science. The Journal of Chemical Physics 149, 180901 (2018).

41. COVID-19 Molecular Structure and Therapeutics Hub. https://covid.molssi.org/.

